# Impact of scaffolding protein TNRC6 paralogs on gene expression and splicing

**DOI:** 10.1101/2021.02.09.430449

**Authors:** Samantha T. Johnson, Yongjun Chu, Jing Liu, David R. Corey

## Abstract

TNRC6 is a scaffolding protein that bridges interactions between small RNAs, argonaute (AGO) protein, and effector proteins to control gene expression. There are three paralogs in mammalian cells, *TNRC6A, TNRC6B, and TNRC6C*. These paralogs have ~40% amino acid sequence identity and the extent of their unique or redundant functions is unclear. Here, we use knockout cell lines, enhanced crosslinking immunoprecipitation (eCLIP), and high-throughput RNA sequencing (RNAseq) to explore the roles of TNRC6 paralogs in RNA-mediated control of gene expression. We find that that the paralogs are largely functionally redundant and changes in levels of gene expression are well-correlated with those observed in *AGO* knockout cell lines. Splicing changes observed in *AGO* knockout cell lines are observed in *TNRC6* knockout cells. These data further define the roles of the TNRC6 isoforms as part of the RNA interference (RNAi) machinery.

## INTRODUCTION

Scaffolding proteins play critical roles in biology by bringing proteins with diverse functions into proximity (Shaw and Filbert 2009). Their ability to guide the formation of complexes increases the effective concentrations of proteins and nucleic acids relative to one another, allowing for more efficient activities inside cells. One important example of scaffolding proteins is the GW182 family (Eystathioy et al. 2002) which plays a critical role facilitating the regulation of gene expression during RNA interference (RNAi) (Baillat and Shiekhattar 2009; Takimoto et al. 2009; Niaz and Hussain 2018).

In vertebrates, there are three GW182 protein paralogs, also known as trinucleotide repeat containing protein 6A (TNRC6A), TNRC6B, and TNRC6C. These are multidomain proteins consisting of an argonaute (AGO) binding domain that can bind up to three AGO protein paralogs (Nishi et al. 2013; Pfaff et al. 2013; Elkayam et al. 2017), CCR4-NOT interacting motif (CIM) (Chekulaeva et al. 2011; Fabian et al. 2011), an ubiquitin associated-like (UBL) domain (Nishi et al. 2013), a glutamine rich domain (Q-rich) (Baillat and Shiekhattar 2009), a PABP-interacting motif 2 (PAM2) (Fabian et al. 2009; Lazzaretti et al. 2009), and an RNA recognition motif (RRM) (Eulalio et al. 2009) (**Figure 1A**).

**FIGURE 1.**
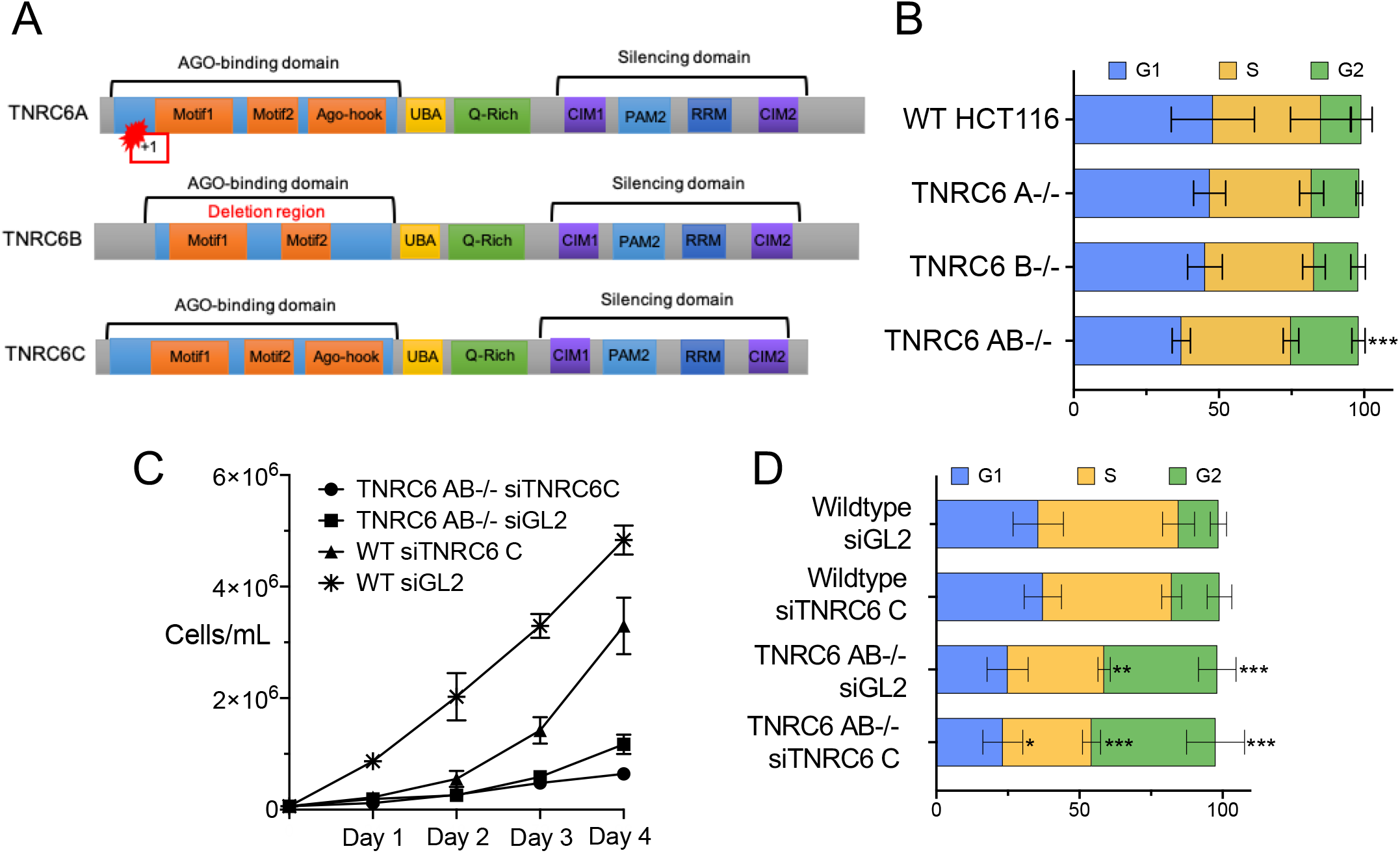
Effect of loss of *TNRC6A, TNRC6B*, or *TNRC6C* expression on cell cycle and cell proliferation. (*A*) Diagram of TNRC6A, TNRC6B, and TNRC6C proteins, with known motifs, knockout mutations, and deletions. (*B*) Percentage of cells in each stage of the cell cycle. (*C*) Growth curve for cell lines transfected with anti-*TNRC6C* siRNA. (*C*) Percentage of cells in each stage of the cell cycle for transfected cell lines wild type after transfection with siTNRC6C, or control duplex siGL2. * p-value > 0.05; ** p-value > 0.01. *** p-value > 0.001.

For RNA interference, TNRC6 plays a critical bridging role. AGO protein binds miRNA guide strands and the miRNA:AGO complex associates with complementary target RNA sequences. TNRC6 binds to AGO through the one of two (in TNRC6B) or three (TNRC6A or TNRC6C) motifs in the AGO binding domain (Elkayam et al. 2017) (**Figure 1A**). The ability of TNRC6 to bind multiple AGO proteins permits enhanced association through cooperative binding between two or three AGO:miRNA complexes (Broderick et al. 2011; Gebert and MacRae 2019; Briskin et al. 2020). Mass spectrometry has identified other accessory proteins binding to TNRC6 domains that may contribute to the control of gene expression (Hicks et al. 2017; Suzawa et al. 2017; Sarshad et al. 2018). The best known TNRC6 interactor is the CCR4-NOT complex that is responsible for translation repression during RNAi (Behm-Ansmant et al. 2006; Fabian et al. 2011; Collart 2016) but was initially discovered as a regulator of gene transcription (Albert et al. 2000).

This partnership between AGO and TNRC6 proteins is central to understanding how RNAi governs gene expression. Here, we use *TNRC6* knockout and knockdown cells deficient in *TNRC6A*, *TNRC6B*, and *TNRC6C* expression in combination with enhanced crosslinking immunoprecipitation (eCLIP) (van Nostrand et al, 2016) and RNA sequencing (RNAseq) to investigate the potential for unique and redundant function for the TNRC6 paralogs during RNAi. We find that the TNRC6 paralogs are largely redundant and that effects on gene expression are remarkably consistent to those observed when AGO proteins are knocked out.

## RESULTS

### Experimental Design: TNRC6 knockout and knockdown cells

We have previously described *TNRC6 A-/-, TNRC6 B-/-, and TNRC6 AB*-/- knockout cell lines (Liu et al. 2019). HCT116 colorectal cancer cells were chosen as a parental line because they are diploid, which facilitates knocking out multiple genes simultaneously. The use of HCT116 as a parental line also allows us to compare the TNRC6 lines directly with AGO knock out cell lines that were also created from HCT116 cells ((Chu et al. 2020), Y Chu, S Yakota, J Liu et al., 2021, accompanying manuscript).

The knockout of the TNRC6A was confirmed by western blot analysis (**Supplementary Figure 1A**). We did not possess an adequate antibody to detect TNRC6B and knockout of TNRCB protein expression was confirmed by mass spectrometry (Liu et al. 2019). The knockout of *TNRC6 A* was due to a point mutation that produced a frameshift while the knockout of *TNRC6 B* was achieved by a 95,481 base-pair deletion (**Figure 1A**).

*TNRC6 A*-/- or *TNRC6 B*-/- single knockout cells grew slower than wild cells (Liu et al. 2019). *TNRC6 AB*-/- double knockout cells were the slowest proliferating. We could not obtain a *TNRC6 ABC*-/- triple mutant, consistent with residual TNRC6 function being necessary for cell growth. To examine the effect of loss of TNRC6 C we used a pool of anti-TNRC6C duplex RNAs that knocked down greater than 90% of *TNRC6C* expression (**Supplementary Figure 1B**).

### Effect of TNRC6 paralog expression on cell cycle

To further evaluate the impact of the TNRC6 paralogs on cell proliferation, we examined the consequences of TNRC6 knockout on cell cycle. The *TNRC6 A*-/- and *TNRC6 B*-/- knockout cell lines showed no significant changes throughout the cell cycle (**Figure 1B**). Double knockout *TNRC6 AB*-/- cells, had a significant increase in G2 phase in the *TNRC6 AB*-/- cells (**Figure 1B**). The change in cell cycle stage for the double knock out cells is consistent with the reduced cell growth seen in the *TNRC6 AB*-/- cells (Liu et al., 2019).

Because we could not obtain triple knockout *TNRC6 ABC*-/- cells, we used a siRNA to knock down TNRC6C expression (**Supplemental Figure 1B**). As a control, we also transfected a noncomplementary duplex RNA, siGL2, into both wild-type HCT116 and *TNRC6 AB*-/- double knockout cells. We observed that cell growth was decreased by transfection of the anti-TNRC6C duplex RNA relative to wild-type cells but that knocking down TNRC6 C has little effect for *TNRC6 AB*-/- double knockout cells (**Figure 1C**).

When TNRC6 C alone is knocked down using an siRNA pool, the cell cycle does not change significantly relative to addition of the control duplex (**Figure 1D**). When both TNRC6A and TNRC6B are knocked out, the addition of either control or anti-TNRC6C duplex RNA in complex with lipid dramatically alters the cell cycle. Transfection with lipid can stress cells. These data suggest that the loss of most function of the three TNRC6 paralogs may have a bigger impact when the cells are challenged by environmental change.

### Impact of TNRC6 knockout/knockdown on gene expression

We used RNA sequencing (RNAseq) of whole cells to evaluate the impact of knocking down or knocking out TNRC6A, TNRC6B, and TNRC6C on overall gene expression (**Figure 2**). Read depth was consistent across all samples. Knocking out TNRC6A had a greater effect on gene expression than knocking out TNRC6B or knocking down TNRC6C (**Figure 2A**). The *TNRC6 AB*-/- double knockout has a larger effect than the single knockout cell lines (**Figure 2A**). The greatest impact on gene expression was observed for the combination of *TNRC6 AB*-/- knockout and TNRC6C siRNA-mediated knockdown (**Figure 2A**).

**FIGURE 2.**
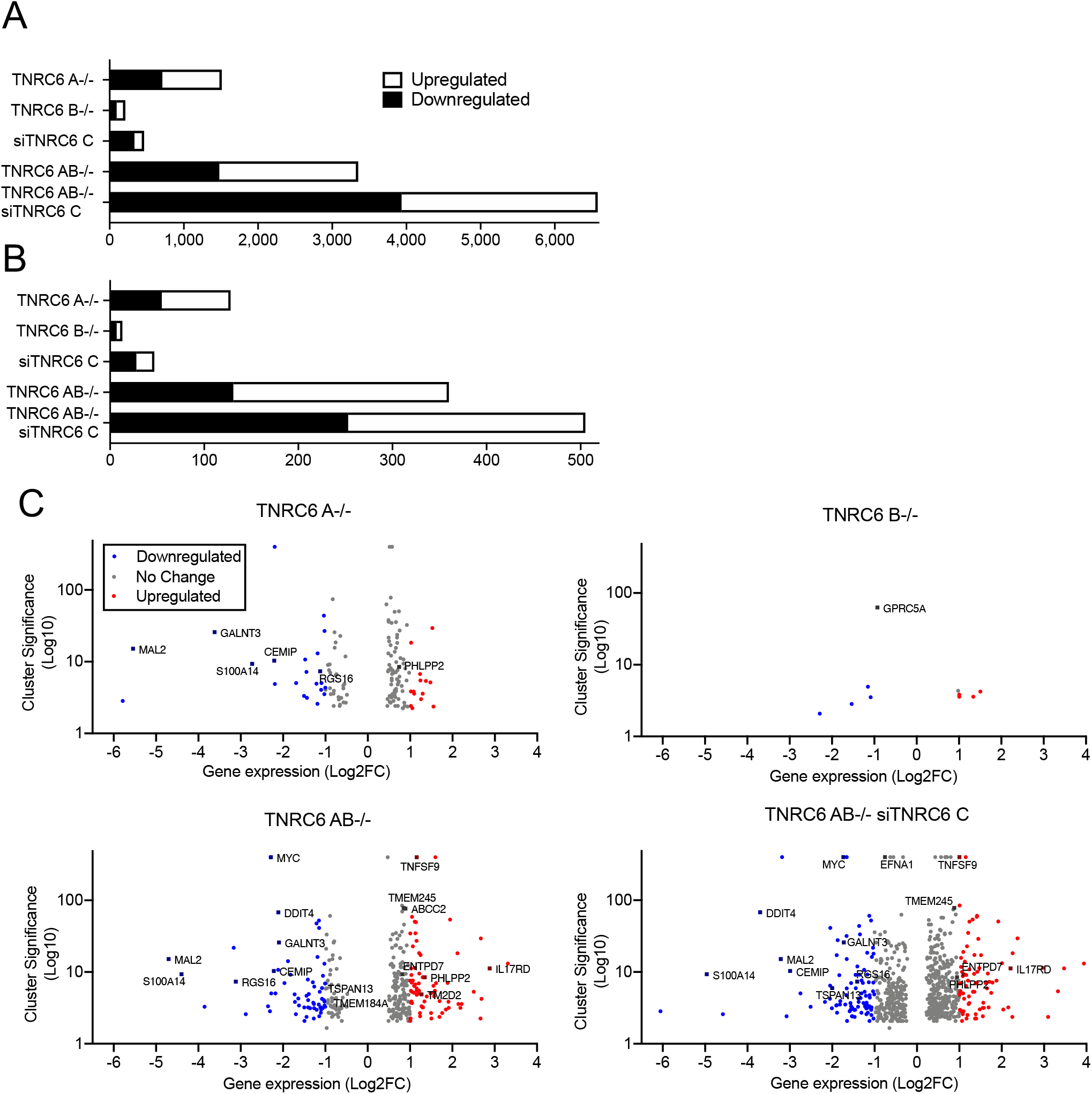
Association of AGO2 protein binding and gene expression in *TNRC6* knockout cells. (*A*) Total number of significantly up-or down-regulated genes in knockout cell lines. (B) Total number of significantly up-or down-regulated genes in knockout cell lines. that overlap with AGO2 binding sites in coding sequences (CDS) and in the 3’ untranslated regions (3’UTR). (*C*) Volcano plots of gene expression in *TNRC6* knockout cell lines.

We have previously used enhanced crosslinking immunoprecipitation (eCLIP) (Van Nostrand et al. 2016) to identify locations within the transcriptome where AGO2 binds (Chu et al., 2020). Genes that host these sites are candidates for regulating gene expression because significant AGO2 binding is thought to be correlated with recognition of miRNAs (Lewis et al. 2003; Friedman et al. 2009; Gebert and MacRae 2019; Chu et al. 2020; Eisen et al. 2020). These regions were identified by clusters of RNAseq reads that were not present or not significantly enriched when compared to parallel experiments using AGO2 knockout cells or a size-matched input sample.

The standard mechanism for endogenous miRNA regulation in mammalian cells suggests that regulation is through interactions within the 3’-untranslated region (3’-UTR). Therefore, we examined mRNAs that possessed read clusters within their 3’-UTRs. We then measured the effect of TNRC6 loss on the expression of these genes in knockout versus wild-type cells (**Figure 2B**).

In all cell lines examined, only a small fraction of gene expression changes (**Figure 2A**) were associated with a significant AGO2 binding clusters (**Figure 2B**). As we had observed for overall gene expression, the number of genes with altered expression was less in single knockout cells, greater in double knockout cells, and greatest in the *TNRC6 AB*-/- siTNRC6C cell line. Once again, the *TNRC6 A*-/- knockout had a bigger effect on expression than the *TNRC6 B*-/- knockout or siTNRC6C knockdown.

A standard assumption of miRNA action is that binding of an AGO:miRNA complex within the 3’-UTR will repress gene expression (Friedman et al. 2009; Guo et al. 2010; Jonas and Izaurralde 2015; Gosline et al. 2016; Gebert and MacRae 2019) and that knocking out AGO variants should increase gene expression. While reducing expression of RNAi factors like the TNRC6 variants would be expected to produce a complex mix of gene expression changes – some direct and some indirect – genes that associate with AGO2 would be expected to be de-repressed when critical RNAi factors are knocked out or knocked down.

We observed, however, that regardless of whether we examine the expression of all genes (**Figure 2A**) or only genes with AGO binding clusters (**Figure 2B**), that similar numbers of genes were associated with up-and down-regulation. These data indicate that there is no simple correlation between AGO2 occupancy and up-or downregulation of a transcript. We then used volcano plots to visualize cluster significance and fold change of individual genes (**Figure 2C**). Once again, the *TNRC6 A*-/- cell line showed more profound changes than *TNRC6 B*-/- cells. The *TNRC6 AB*-/- cells or *TNRC6 AB*-/- siC cells showed much greater effects than the single gene knockout cells, both in terms of the number of genes changed and the magnitude of gene expression changes. Cluster significance, the indication of AGO2 occupancy, was not associated primarily with up or down regulation regardless of which knockout cell line is examined.

### Comparing the impact of AGO and TNRC6 knockouts on global gene expression

The TNRC6 paralogs are important binding partners for AGO proteins (Kalantari et al. 2016b; Hicks et al. 2017). The scaffolding domains of TNRC6 facilitate the recruitment of effector proteins for mRNA degradation (Piao et al. 2010; Chekulaeva et al. 2011; Fabian et al. 2011; Hicks et al. 2017) or transcriptional activation (Hicks et al. 2017; Liu et al. 2018; Liu et al. 2019). Because of the partnership between AGO and TNRC6 we hypothesized that many gene expression changes would be shared between *TNRC6* and *AGO* knockout cells and be the best candidates as endogenous control points for regulation by miRNAs. It is also possible, however, that AGO and TNRC6 proteins may play independent roles. To evaluate these hypotheses and identify candidate genes, we compared the impact on gene expression of knocking out AGO and TNRC6 proteins (**Figure 3**).

**FIGURE 3.**
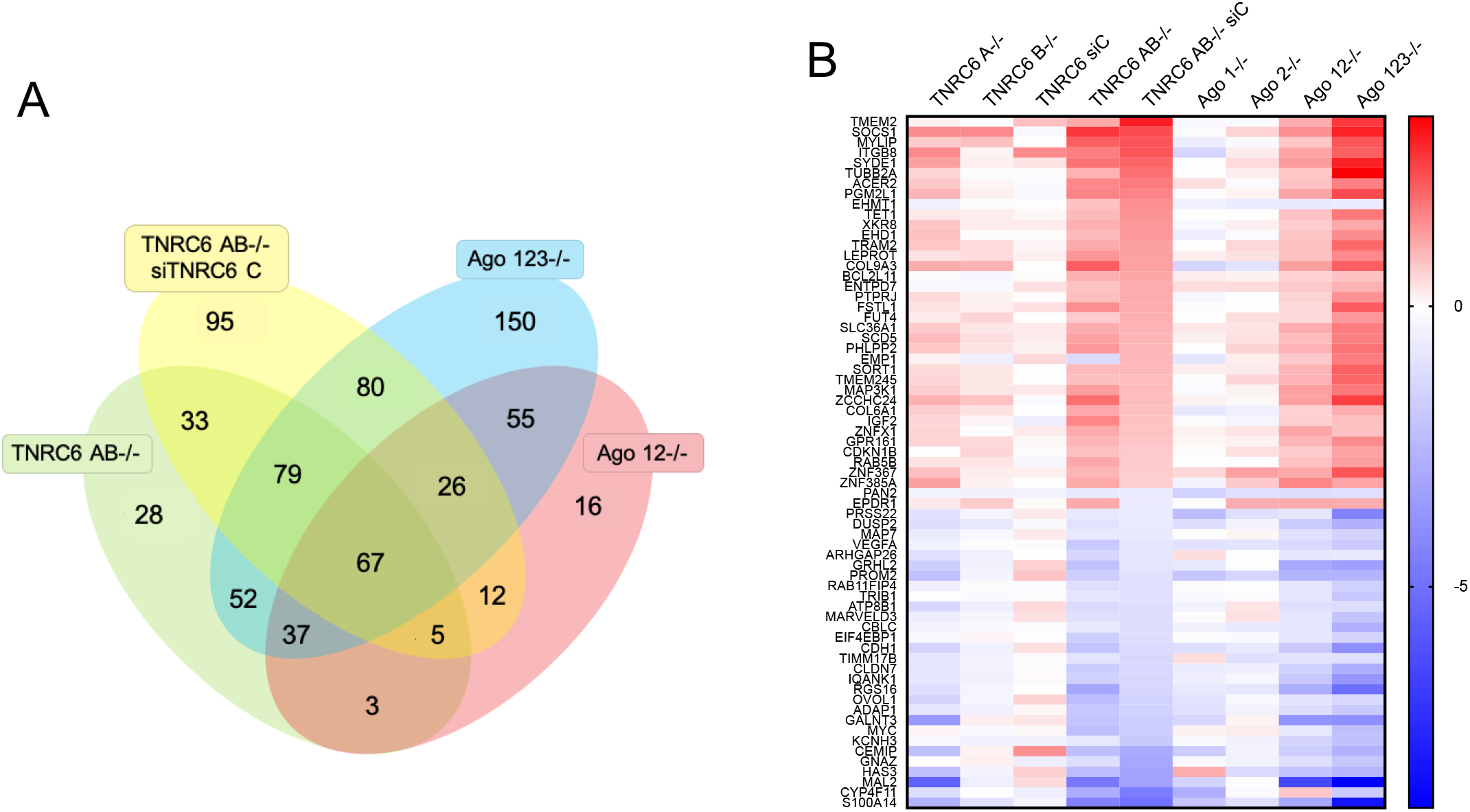
Consistent variation for gene expression changes in *TNRC6* and *AGO* knockout cells. (*A*) Venn diagram showing the overlap of gene expression changes associated with AGO2 binding within 3’-UTRs. (*B*) Heatmap showing gene expression changes shared by *TNRC6* and *AGO* knockout cell lines.

We focused on genes that had significant AGO2-binding clusters within their 3’-UTRs (Chu et al. 2020) and compared the gene expression changes in *TNRC6 AB-/-, TNRC6 AB*-/- siTNRC6C, *AGO12*-/-, and *AGO123*-/- cells relative to wild-type HCT116 cells (**Figure 3A**). When examining large datasets, it is important to prioritize outputs. We reasoned genes that genes showing expression changes in multiple cell lines would be the best candidates for physiologically relevant gene regulation. Gene expression changes due to experimental noise or artefacts from RNAseq are least likely when the changes occur in multiple cell lines. We recognize that these stringent criteria may overlook some candidates, but they facilitate focusing on a manageable number of genes for further analysis.

We identified sixty-seven genes with AGO2 binding clusters and significantly changed gene expression (FDR<0.05, −0.6>Log_2_ Fold Change>0.6) that were shared in all of the four cell lines (**Figure 3A**). These sixty-seven genes included examples of both up-and down-regulation, with thirty-six genes increasing expression and thirty-one genes with reduced expression.

We then used heat map analysis to sort these genes according to altered gene expression and to extend the comparison to our *AGO* knockout cell lines. (**Figure 3B**). Of the nine cell lines examined, the siTNRC6C knockdown cells showed the least change and no obvious correlation for up-or down-regulation. Of the remaining *TNRC6* knockout cell lines, gene expression changes were weakest in *TNRC6 B*-/- cells, stronger in *TNRC6 A*-/- cells, and strongest in the *TNRC6 AB*-/- and *TNRC6 AB*-/- siC cells – a comparison reminiscent of our data for cell proliferation, cell cycle analysis, and global gene expression. For the eight knockout cell lines, genes that were upregulated in *AGO* knockout cells tended to be upregulated in *TNRC6* knockout cells, while genes that were downregulated in *AGO* knockout cells showed similar downregulation in the engineered *TNRC6* cells (**Figure 3B**).

For comparison, we also examined gene expression changes in other overlapping cohorts of knockout cells. For example, the 95 genes that were only changed significantly in the *TNRC6 AB*-/- siTNRC6C cell lines (**Figure 3A**), did not show similar gene expression trends relative to the other cell types (**Figure 4A**). This result supports the conclusion that these gene expression changes are unrelated to perturbation of the RNAi pathway. Conversely, the 185 changes shared between the *AGO123*-/- and the *TNRC6 AB*-/- siTNRC6C mostly overlap (**Figure 4B**), suggesting that these genes are more likely to be regulated by RNAi (**Figure 4B**).

**FIGURE 4.**
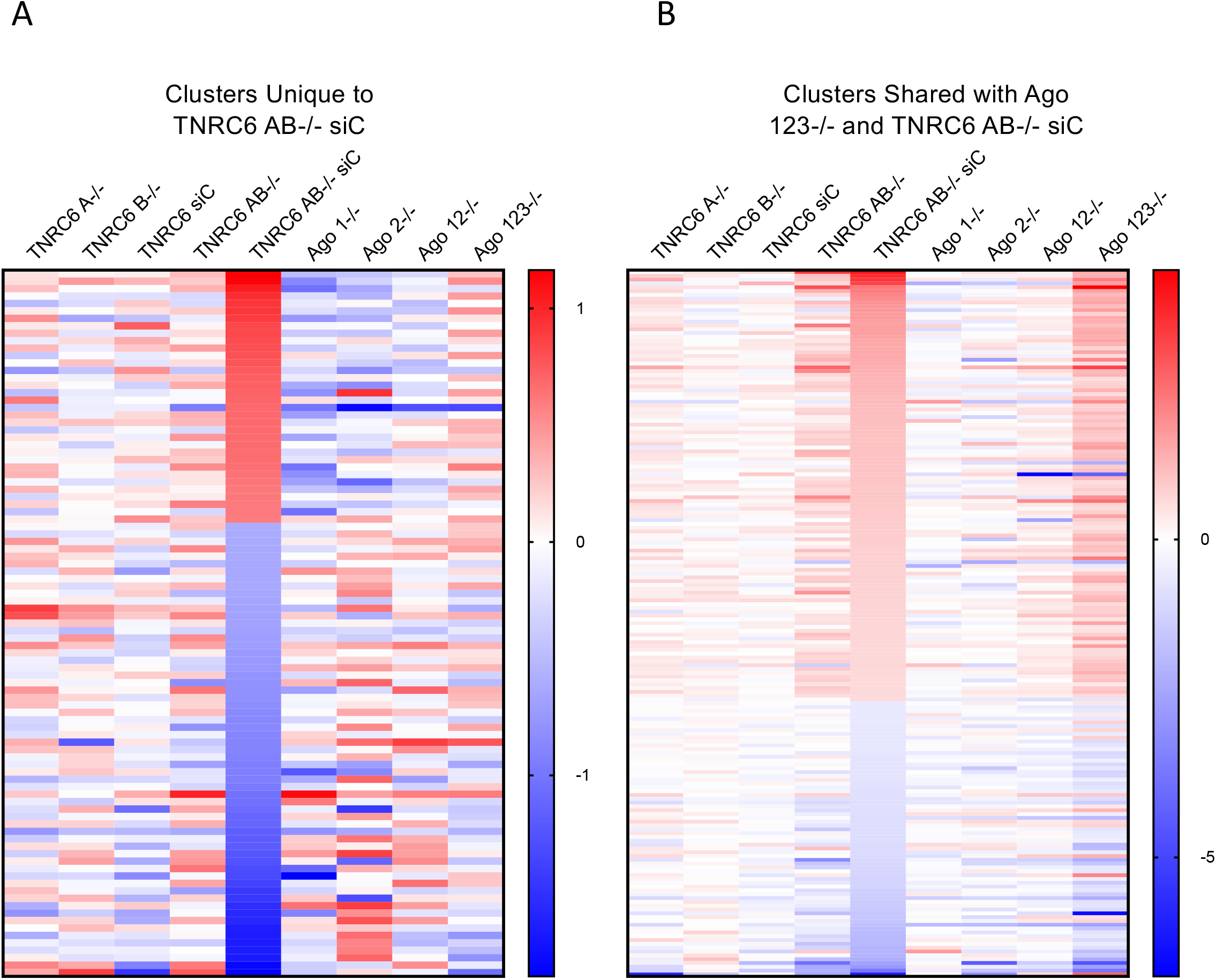
Comparison of gene expression in *TNRC6* and *AGO* knockout cells**. (**A) Heatmap of gene expression changes of 95 genes with AGO2 protein binding clusters that change significantly only in the *TNRC6AB*-/- knock out/si*TNRC6C* knockdown cells. (B) Heatmap of gene expression changes of 159 genes with AGO2 clusters that change significantly in *TNRC6 AB-/-siTNRC6C* and *AGO123*-/- cells.

When evaluating CLIP-seq data, it is essential to view the primary data to evaluate the characteristics of each cluster of reads to ensure the quality of the data and identify different classes of read cluster. Previously, we had focused our experimental validation of gene expression in AGO knockout cells on 22 representative genes with AGO2-binding clusters (Chu et al. 2020). These genes were chosen to represent differing species of highly significant cluster (single clusters versus multiple closely space clusters), and both up-and down-regulated genes. These cluster sites contained seed sequence complementary to well-expressed miRNAs.

We compared the gene expression of these twenty-two chosen genes in our nine knockout or knockout/knockdown datasets. Similar to the result observed with our sixty-seven gene overlapping cohort (**Figure 4**), the selected twenty-two genes showed a similar rank order of gene expression change regardless of whether AGO or TNRC6 variants were being knocked out (**Figure 5A**). As observed for the sixty-seven gene cohort, the siTNRC6C knockdown cells did not trend with the other cell lines, suggesting that the TNRC6C knockdown had the least impact on cells. These data demonstrate that the broad trends correlating the impact of gene expression of TNRC6 and AGO knockdown also apply to genes with strongest AGO2 association detected by eCLIP.

**FIGURE 5:**
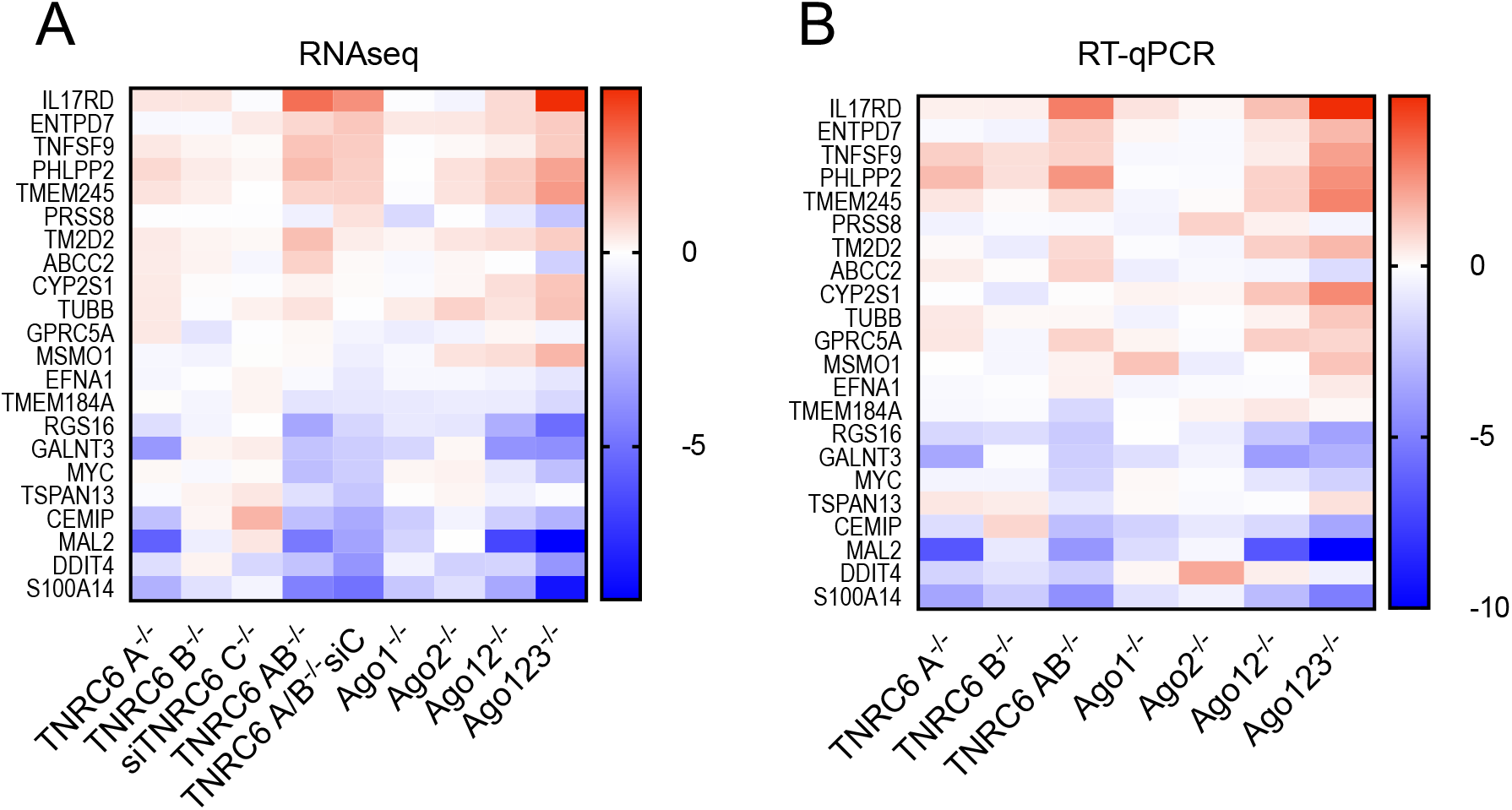
Consistent variation of gene expression in cells with highly ranked AGO2 protein binding clusters. (A) Heatmap of gene expression changes from *TNRC6* knockout and *AGO* knockout cell lines from RNA sequencing for 22 cluster genes examined in Chu et al, 2020. (B) Heatmap of gene expression changes from *TNRC6* knockout and *AGO* knockout cell lines from RT-qPCR for 22 cluster genes examined in Chu et al, 2020.

Quantitative PCR (qPCR) was performed to validate the RNAseq data (**Figure 5B**). Measurement of RNA samples from each cell line confirmed the trends observed the RNAseq data. These data suggest that the trends correlating *AGO* or *TNRC6* knockout remain similar regardless of the shape of the RNA read cluster detected by eCLIP and RNAseq.

### Impact of AGO and TNRC6 variants on alternative splicing

RNAi has been suggested to have the potential to directly regulate splicing (Allo et al. 2009; Liu et al. 2012; Liu et al. 2015; Fuchs et al. 2021). In a related study, we have examined the impact of knocking out AGO variants on gene splicing (Y Chu, S Yakota, J Liu et al., 2021, accompanying manuscript). We now examine the impact of knocking out TNRC6 paralogs on splicing to assess the involvement of TNRC6 on the regulation of endogenous splicing by miRNAs.

We evaluated the changes in splicing observed in our knockout cell lines. Venn diagrams were used to visualize all skipped exon splicing events that were changed in *AGO123-/-, AGO12-/-, TNRC6AB*-/-, and *TNRC6AB*-/- siTNRC6C cell lines (**Figure 6AB**). Changes that were observed in all four lines were awarded the highest priority for analysis because we reasoned that shared events would be most likely to have physiological relevance.

**FIGURE 6.**
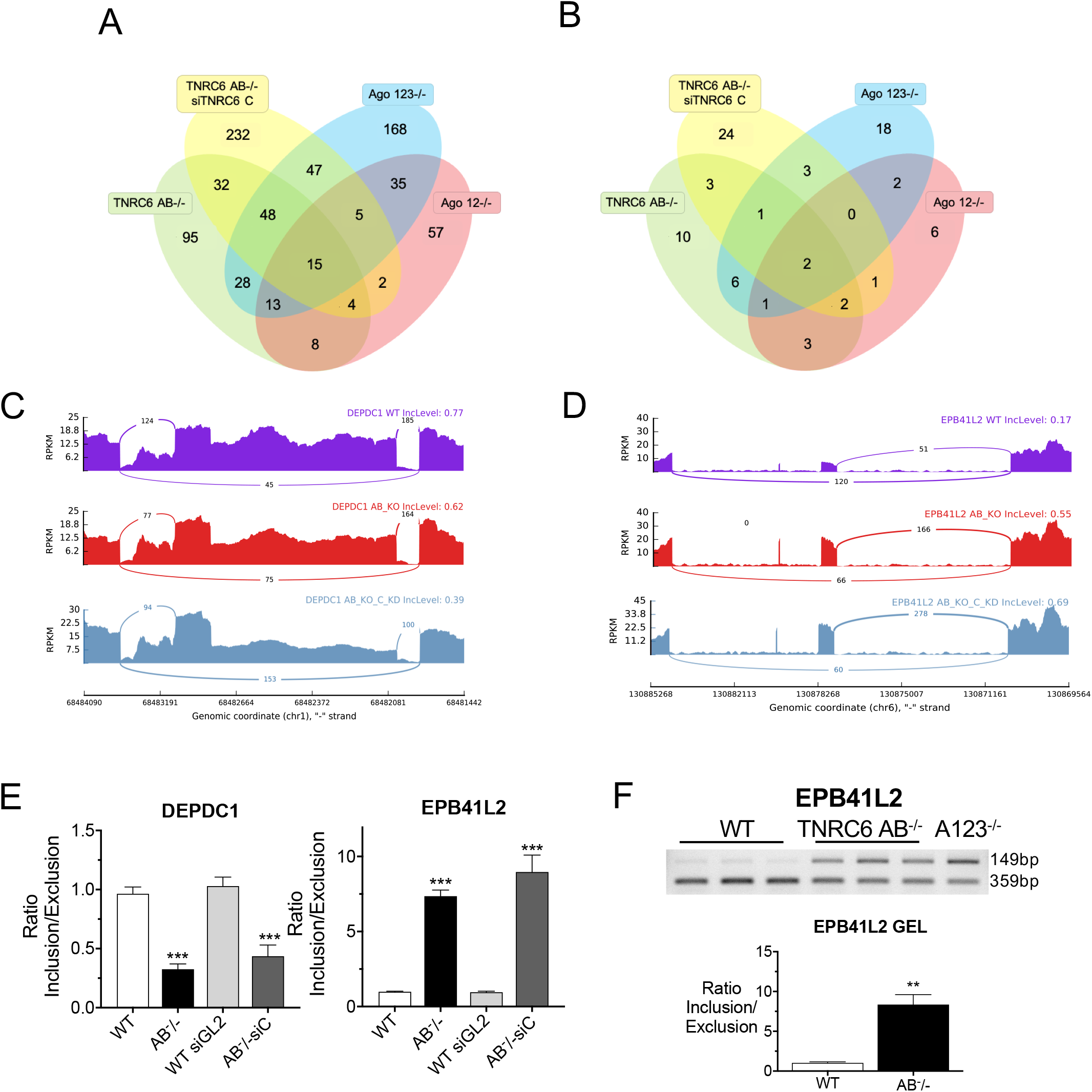
Changes in alternative splicing in *TNRC6* knockout cell lines. (*A*) Venn diagram of skipped exon splicing events. (*B*) Venn diagram of skipped exon splicing events located near AGO2 binding clusters. (*C*+*D*) sashimi plots for genes *DEPDC1* and *EPB41L2* that overlap between *TNRC6 AB*-/- and *TNRC6 AB-/-siC* RNAseq datasets in (*B*). (E) qPCR validation of skipped exon events in *TNRC6 A/B* knockout and *TNRC6 A/BKO/siC* cells. Error bars represent standard deviation (SD). *P < 0.05; **P < 0.01; ***P < 0.001 compared with control cell by two tailed *t*-test. (F) Semiquantitative PCR validation of skipped exon events in *TNRC6 A/B* KO cells and quantitation of the gel.

We found that fifteen skipped exon splicing events are shared between the four cell lines (**Figure 6A**). Of those fifteen, only two genes *(DEPDC1* and *EPB41L2)* had significant AGO2 binding clusters located near affected introns (**Figure 6B).** Visual inspection of the sashimi plots for *DEPDC1* showed an increase in exon skipping in the *TNRC6 AB*-/- and *TNRC6 AB*-/- siC cell lines relative to wildtype **(Figure 6C**) while *EPB41L2* sashimi plots showed a decrease in exon skipping for the knockout cell lines (**Figure 6D**). The splicing changes seen in the RNAseq data for *DEPDC1* and *EPB41L* were validated by qPCR (**Figure 6E**). Splicing changes for *EPB41L2* were further validated by PCR gel (**Supplementary Figure 3**).

Our criteria that changes occur in all cell lines is restrictive. Intronic RNA is recovered at relatively low amounts and we recognized that our criteria might cause us to overlook some candidates. We chose, therefore, to examine the impact of TNRC6 knockouts on seven genes that our laboratory had already evaluated for splicing changes due to knockout of AGO proteins (Y Chu, S Yakota, J Liu et al., 2021, accompanying manuscript) (**Figure 7**). These seven genes were chosen because they had at least one AGO2 cluster near a significant splicing event locus within an intron and had a miRNA candidate complementary to a sequence within the AGO2 cluster.

**FIGURE 7.**
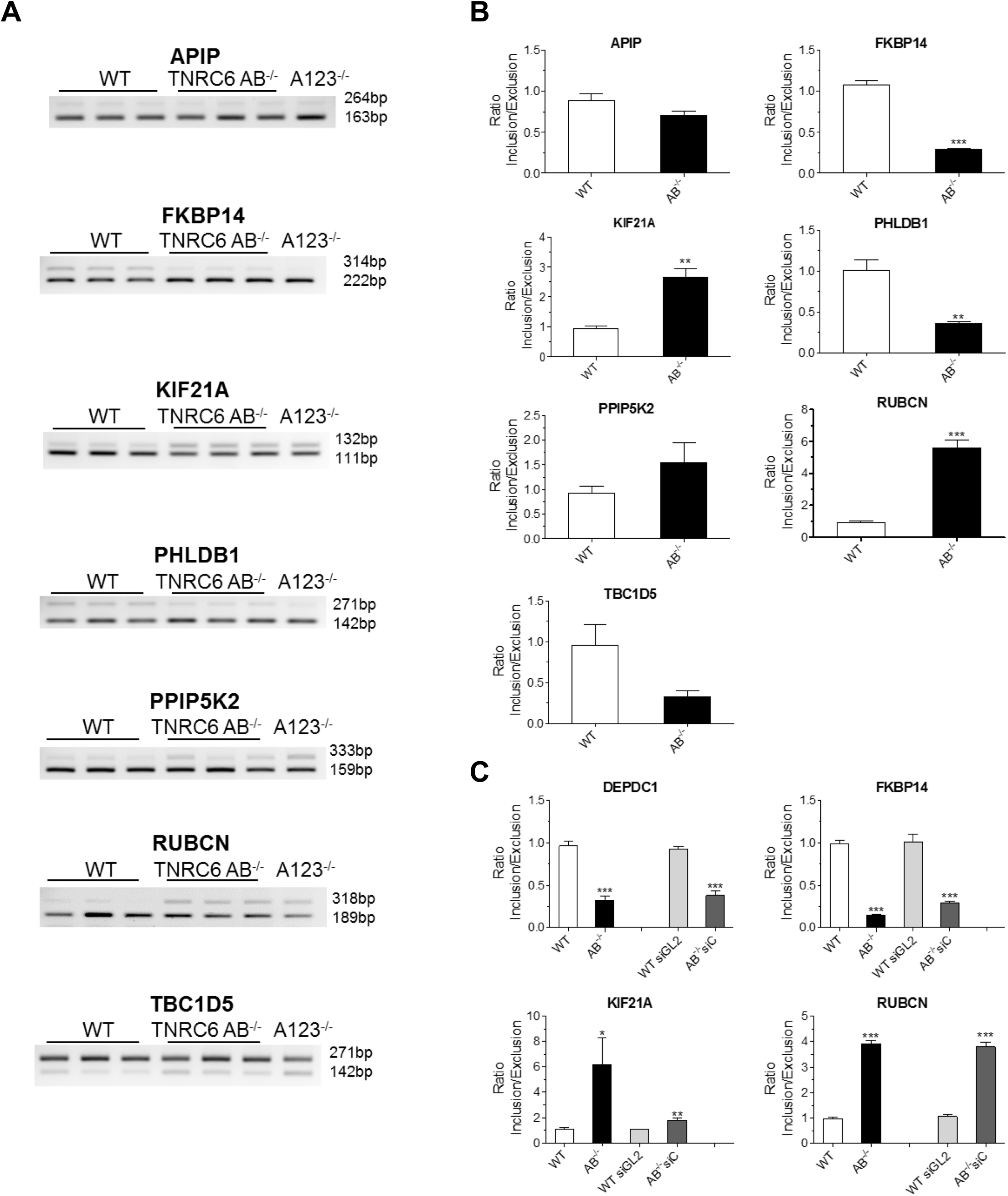
Validating the effect of *TNRC6* knockouts on alternative splicing. (*A*) Semiquantitative PCR validation of skipped exon events in *TNRC6 A/B* knockout cells. (*B*) Quantitation of data shown in in part (A). (*C*) QPCR validation of skipped exon events in *TNRC6 A/B* knockout and *TNRC6 A/B knockout/siCTNRC6* knockdown cells. Error bars represent standard deviation (SD). *P < 0.05; **P < 0.01; ***P < 0.001 compared with control cell by two tailed *t*-test.

Of the seven genes analyzed, our RNAseq data for TNRC6 knockout cells showed significant changes in alternative splicing for five genes in either the *TNRC6 AB*-/- or *TNRC6 AB*-/- siC cells. We evaluated splicing changes by reverse transcriptase PCR (**Figure 7AB**). For five genes, *FKBP14, KIF21A, PHLDB1, RUBCN*, and *TBC1D5*, RT-PCR data confirmed significant splicing changes in both *TNRC6 AB*-/- and *AGO 123*-/- cells. (**Figure 7B**). For *FKBP14* and *KIF21A*, qPCR data confirmed splicing changes in *TNRC6 AB*-/- and *TNRC6 AB*-/- siC (**Supplementary Figure 4**). Two genes, *APIP* and *PPIP5K2*, did not have significant detected changes in RNAseq data and we also did not observe significant changes in PCR (**Figure 7AB**).

Two genes, *RUBCN* and *FKBP14*, that showed splicing changes in the TNRC6 knockout cells were examined in more detail in an accompanying study (Y Chu, S Yakota, J Liu et al., 2021, accompanying manuscript). Their splicing was shown to be modulated by miRNA mimics or anti-miRNA oligonucleotides that target the site for AGO2 association determined through eCLIP. These data reinforce the conclusion the suggestion that the genes are targets for endogenous small RNAs.

## DISCUSSION

### TNRC6 paralogs and RNAi

The three human TNRC6 paralogs, TNRC6A, TNRC6B, and TNRC6C, play important roles in RNAi (Baillat and Shiekhattar 2009; Niaz and Hussain 2018). The complex between a small RNA and AGO proteins recognizes the target sequences within cellular RNA, while the three multi-domain TNRC6 paralogs bind to AGO and act as scaffolds to promote association with proteins that modulate function. While the three protein paralogs are ~40% identical, their potential for unique or redundant activities has not been determined. Neither has the extent to which their impact on endogenous gene expression overlaps the impact of AGO proteins.

We had previously investigated the impact of knocking out TNRC6 paralogs on the ability of synthetic duplex RNAs to control translation and splicing (Liu et al., 2019). In those studies, we found that knocking out TNRC6 expression did not affect inhibition of translation or splicing by fully or highly complementary synthetic duplex RNAs but did reverse the action of synthetic miRNA mimics. Here we use knockout cell lines and an efficient siRNA pool that reduces TNRC6C expression to analyze the role of TNRC6 expression on global gene expression.

### Redundancy or independence: Roles for TNRC6 paralogs

We had previously used knockout cells to demonstrate that the loss of both TNRC6A and TNRC6B was required to affect gene silencing by synthetic duplex RNAs (Liu et al. 2019). Alone, TNRC6A and TNRC6B were replaceable. Consistent with this observation, we now observe knocking out TNRC6A or TNRC6B alone has no significant effect on the cell cycle (**Figure 1**). Larger impacts on cell cycle were observed in the *TNRC6 AB*-/- double knockout cell line or *TNRC6 AB*-/- siTNRC6C cells (**Figure 2**). Analysis of gene expression or alternative splicing revealed similar results. Blocking expression of two or three TNRC6 paralogs affected expression (**Figure 3**) or alternative splicing (**Figure 5**) of many more genes than did blocking expression of TNRC6A, TNRC6B, or TNRC6C alone. The conclusion that TNRC6 paralogs have largely redundant functions regulating endogenous gene expression is consistent with our previous observation of redundant function when modulating the effects of designed synthetic RNAs.

While the single knockouts showed relatively little change relative to double knockout cells, we did observe substantially larger number of genes with altered expression in the *TNRC6 A*-/- cells than in *TNRC6 B*-/- cells. These data may indicate that that TNRC6A plays a unique role in regulating expression of a subset of genes. Alternatively, *TNRC6A* is the most highly expressed paralog (**Supplementary Figure 2**) and its loss from the pool of TNRC6 protein may have the biggest impact.

### How do TNRC6 paralogs affect regulation by RNAi?

We have used eCLIP to identify sites for AGO2 binding within cytoplasmic (Chu et al., 2020) and nuclear (Y Chu, S Yakota, J Liu et al., 2021, accompanying manuscript). RNA and used these data to understand how AGO binding correlates with gene expression and splicing at these sites. One conclusion from these studies was that, contrary to the standard expectation that AGO2 binding within a 3’-UTR should be associated with gene repression, we observed that genes with significant association to AGO2 showed by up-and down-regulation upon gene knockouts.

Here we report that the effects of knocking out TNRC6 paralogs on the expression of genes with significant AGO2 binding sites yield remarkably similar results (**Figure 3**). We can make several conclusions from these data: 1) TNRC6 proteins are largely redundant, although knockout of TNRC6 A has the largest effect. As additional TNRC6 paralogs are removed, effects become greater; 2) The similarity of up-and down-regulated genes reveals the remarkable extent to which AGO and TNRC6 proteins function as partners to control gene expression; 3) As with AGO knockout cells, knocking out TNRC6 paralogs has an unpredictable effect on gene expression. The fact that AGO2 has a significant association with a 3’-UTR cannot be assumed to lead to gene up-regulation when AGO or TNRC6 proteins are removed from cells

We have not resolved whether the changes in gene expression we observe are due to direct effects of miRNAs binding to sites where AGO2 association is detected or indirect effects. Because of the lack of predictable correlation with gene repression or up-regulation, such studies are not straightforward and will be a subsequent focus of research. It is clear from the data, however, that the genes we identify are being controlled by a common RNAi axis that requires expression of both AGO and TNRC6. The fact that we observe both increased and decreased expression at genes with AGO2 association within their 3’-untranslated regions supports our previous conclusion from AGO knockout cells that AGO2 occupancy is not sufficient to infer repression of a target transcript and emphasize the complexity of RNAi function.

### Impact of TNRC6 on alternative splicing

While RNAi is often assumed to be a cytoplasmic mechanism in mammalian cells (Zeng and Cullen 2002), RNAi protein factors and miRNAs also exist in cell nuclei (Gagnon et al. 2014). Functional evidence showing robust control of transcription and splicing by synthetic duplex RNAs (Allo et al. 2009; Liu et al. 2012; Liu et al. 2015; Kalantari et al. 2016a) has suggested the potential for nuclear RNAi to be a natural regulatory mechanism, but persuasive experimental evidence for control of endogenous transcription or spicing has been elusive. In addition, previous studies using synthetic RNAs had shown that TNRC6 is not required for highly complementary synthetic small RNA to influence differential splicing (Liu et al. 2018).

In an accompanying paper, we investigate the impact of endogenous miRNAs and RNAi on alternative splicing (Y Chu, S Yakota, J Liu et al., 2021, accompanying manuscript). We identify sites of AGO2 binding using the same eCLIP dataset used here and correlate AGO2 binding with changes in splicing upon knocking out AGO1, AGO2, and AGO3. We observe changes in splicing and show that splicing can be manipulated by synthetic miRNAs or anti-miRs designed based on predictions of miRNAs that target sites identified by eCLIP.

We now show that knocking TNRC6 variants also affect alternative splicing. As with our AGO datasets, only a relatively small number of candidate splicing events are identified. However, of this small number, there was substantial overlap between our TNRC6 and AGO data, supporting belief they are due to a common RNAi-related pathway. The expression of two of the genes with splicing changes in both *AGO*-/- and *TNRC6*-/- datasets, *RUBCN* and *FKBP14*, could be manipulated by miRNA mimics or antimiRs designed to predicted target sites.

These data suggest that miRNAs have the potential to control endogenous splicing. We had previously reported that duplex RNAs that control splicing did not require expression of TNRC6 (Liu et al., 2019). These RNAs, however, were either fully or almost fully complementary to their target sites within intronic RNA. The scaffolding function of TNRC6 acts to bridge AGO proteins, increasing the cooperativity of binding and allowing imperfectly paired miRNAs to associate with target sequences more tightly (Elkayam et al. 2017). It is possible that, while TNRC6 is not necessary for recognition of highly complementary duplex RNAs, it is necessary for the activity of mixtures of imperfectly complementary miRNAs that act in concert to control endogenous gene expression.

Our data also suggest that AGO and TNRC6 affect the splicing of a relatively small subset of proteins. This outcome is consistent with the conclusion that RNA-mediated regulation of splicing is a minor regulatory mechanism in HCT116 cells. RNAi-mediated regulation of splicing may also be more pervasive in other cell types, cells grown under more demanding environment conditions, during cell development, or during cells involved in disease. Alternatively, intronic RNA is present at relatively low steady state levels (Clement et al. 1999; Mortazavi et al. 2008) and therefore less detectable. Because of our stringent conditions for identifying candidates for AGO2 binding and splicing change, we may be overlooking genes where RNA-directed modulation of splicing is biologically significant yet undetected by our approach.

### Conclusions

Scaffolding proteins play complex roles bringing other proteins, RNA, or DNA together. The TNRC6 family proteins a multi-domain model for understanding the potential of scaffolding proteins to organize molecular function. (Shaw and Filbert 2009) We observe a strong overlap between the changes in gene expression upon knockout of TNRC6 or AGO proteins, consistent with the partnership of these proteins during RNAi. In our HCT116 model cell line, at least, that partnership does not produce predictable changes in gene expression at sites within target RNAs where AGO2 binds. It is clear from our data that TNRC6 plays an important role in controlling gene expression, but there remains much to learn about how it acts in concert with AGO2 and the possibility that it may play significant roles independent of AGO2.

## MATERIALS AND METHODS

### Cell Culture

Wild-type HCT116 cells were obtained from Horizon Discovery. HCT116 cells containing knock out modifications to the *TNRC6A, TNRC6B*, and *TNRC6A & TNRCB* genes were purchased from GenScript. All cell lines were cultured in McCoy’s 5A medium (Sigma-Aldrich) supplemented with 10% FBS (Sigma-Aldrich) in 37°C 5% CO_2_. For cell counting, cells were mixed together with equal volume of trypan blue (Sigma) and were counted using cell counter (TC20^™^ Automated Cell Counter; Bio-Rad).

### Transfections

All transfections used Lipofectamine RNAi MAX (Invitrogen). For transfections, cells were seeded into six-well plates at 150 thousand cells per well for wild type, *TNRC6 A*-/- and *TNRC6 B-/-. TNRC6 AB*-/- cells were seeded at 250 thousand cells per well due to the slowed growth rate of these cells. Cells were transfected as described in Liu et al., 2019.

### PI Staining and FACS

Cells were harvested at 90% confluency for cell cycle analysis by propidium iodide (PI) staining. Cells were harvested using 1X Trypsin-EDTA (Sigma) and washed with PBS. To fix cells, they were suspended in PBS at a concentration of 2×10^6^ cells per mL and added to an equal volume of 100% ethanol while vortexing. Cells were then incubated at −20°C for 24 hours to 1 month. To prepare cells for staining with PI, they were washed three times with PBS to ensure all ethanol was removed. Cells were suspended in staining buffer (0.1% Triton X, 20 ug/mL PI, and 20 μg/mL RNAse A) at 2×10^6^ cells per mL. Stained cells are incubated at 37°C for 15 mins. Cell were then stored at 4°C and protected from the light. Cells were run within 48 hours at the UTSW Flow Cytometry Facility on a Caliber. Data was then analyzed using FlowJo software.

### RNA extraction, RNA sequencing, & enhanced crosslinking immunoprecipitation sequencing (eCLIP)

Whole cell RNA was extracted from cells harvested with trypsin at 90% confluency. RNeasy kit (Qiagen) was used purify the RNA for whole cell steady state mRNA sequencing. RNA sequencing was performed by the McDermott Center Next Generation Sequencing Core at UTSW as described in Chu et al, 2021. RNA sequencing data was analyzed in the same manner as Chu et al. 2021. Methods and analysis for eCLIP were described in Chu et al., 2020.

### qPCR and western blot

Both qPCRs and western blots were performed as described in Liu et al., 2019.

### Splicing analysis by gel electrophoresis and qPCR

Total RNA was extracted from HCT116 wild-type, TNRC6 knockout cells, and treated with DNase I (Worthington Biochemical) at 25 °C for 20 min, 75 °C for 10 min. Reverse transcription was performed using high-capacity reverse transcription kit (Applied Biosystems) per the manufacturer’s protocol. 2.0 μg of total RNA was used per 20 μl of reaction mixture.

For gel electrophoresis analysis, PCR amplification was performed as following; 95 °C 3 min and 95 °C 30 s, 60 °C 40 s, 72 °C 30 s for 35 cycles. The PCR products were separated by 1.5% agarose gel electrophoresis. The bands were quantified by using ImageJ software.

In qPCR analysis for splicing changes by using double strand RNAs and miRNA mimics, PCR was performed on a Biorad CFX384 Real-Time System using iTaq SYBR Green Supermix (BioRad). PCR reactions were done in triplicates at 55 °C 2 min, 95 °C 3 min and 95 °C 20 s, 60 °C 45s for 40 cycles in an optical 384-well plate. The expression level was compared between exon included spliceform and exon excluded spliceform. PCR primers were shown in Supplementary Table 3.

### mRNA quantification

RNA standards of *TNRC6 A, TNRC6 B, TNRC6 C, GAPDH*, and *MYC* synthesized by in vitro transcription from cDNA using SP6 Megascript Kit (ThermoFisher). Primers were designed following the manufactured instructions and are listed in Supplementary Table 1. RNA purity was measured by Bioanalyzer (Agilent, Santa Clara, CA, USA). Purified standard RNA was serially diluted to 100, 1×10^3^, 1×10^4^, 1×10^5^, 1×10^6^, 1×10^7^, 1×10^8^ and 1×10^9^ and used to construct a standard curve for qPCR efficiency following reverse transcription.

Experimental RNA for mRNA quantification was extracted from disassociated and counted cells with seven repeated Trizol/chloroform precipitations. For both the experimental and RNA standard, 0.5 ml DNA LoBind Tubes (Eppendorf) coated in nuclease free water supplemented with RNase Inhibitor and salmon sperm DNA were used to prevent loss of RNA to tube binding.

Total RNA from HCT116 WT, *TNRC6 A-/-, TNRC6 B*-/-, and *TNRC6 AB*-/- was treated with DNase I (Worthington Biochemical) at 25°C for 20 min, 75°C for 10 min. Reverse transcription was performed using high-capacity reverse transcription kit (Applied Biosystems) per the manufacturer’s protocol. PCR was performed on a Biorad CFX384 Real-Time System using iTaq SYBR Green Supermix (BioRad). PCR reactions were done in triplicates at 55 °C 2 min, 95 °C 3 min and 95 °C 20 s, 60 °C 45s for 40 cycles in an optical 384-well plate. The CT values were plotted against the number of molecules (log10-scale) and analyzed with linear regression to calculate the copy number per cell.

## Supporting information

SUPFILES

## DATA AVAILABILITY

All high-throughput sequencing data generated for this study (RNAseq, eCLIP) have been deposited in Gene Expression Omnibus under accession number GSE162749.

## SUPPLEMENTAL MATERIAL

Supplemental material is available for this article.

## ACKNOWLEDGEMENTS

The authors thank Dr. Joshua Mendell for the gift of HCT116:AGO2 knock out cells and Dr. Jay Nelson for anti-AGO antibody 3148. DRC was supported by the National Institutes of Health (NIH) (GM106151) and the Robert Welch Foundation (I-1244). DRC holds the Rusty Kelley Professorship in Medical Science. STJ is supported by an NIH predoctoral fellowship (5 F31 EY030336-03).

## Author contributions

D.R.C. wrote the manuscript and supervised the experiments. S.T.J., Y.C., and J.L. performed the experiments and assisted in writing the manuscript.

