## Supplementary material for "Impact of scaffolding protein TNRC6 paralogs on gene expression and splicing": SUPFILES

A

B

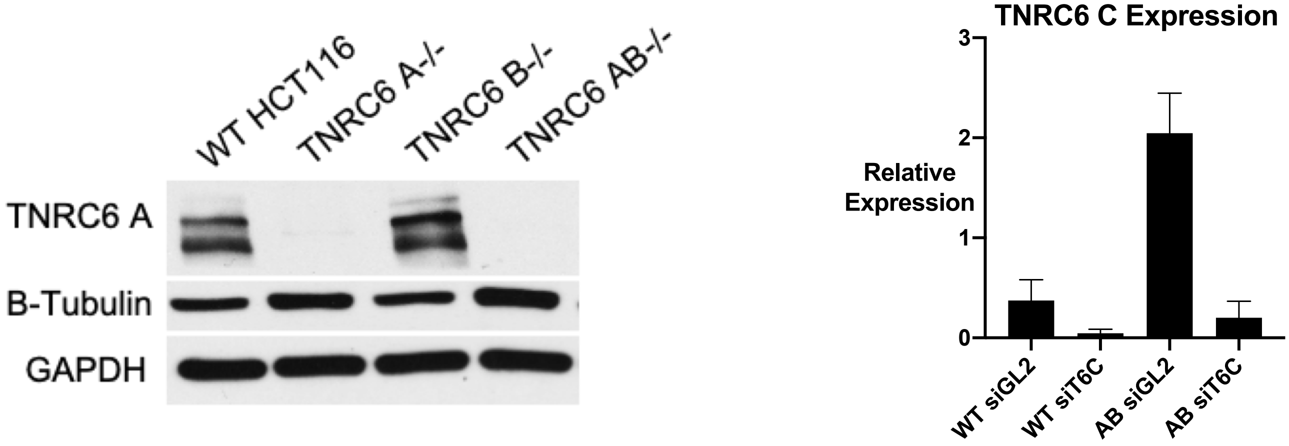


**SUPPLEMENTARY FIGURE 1.** Changes in TNRC6 mRNA and Protein levels after knockout. **(**A) Western blot of TNRC6 A in call cell lines. (B) Bar graph of TNRC6 C relative expression to HPRT in transfected cells.

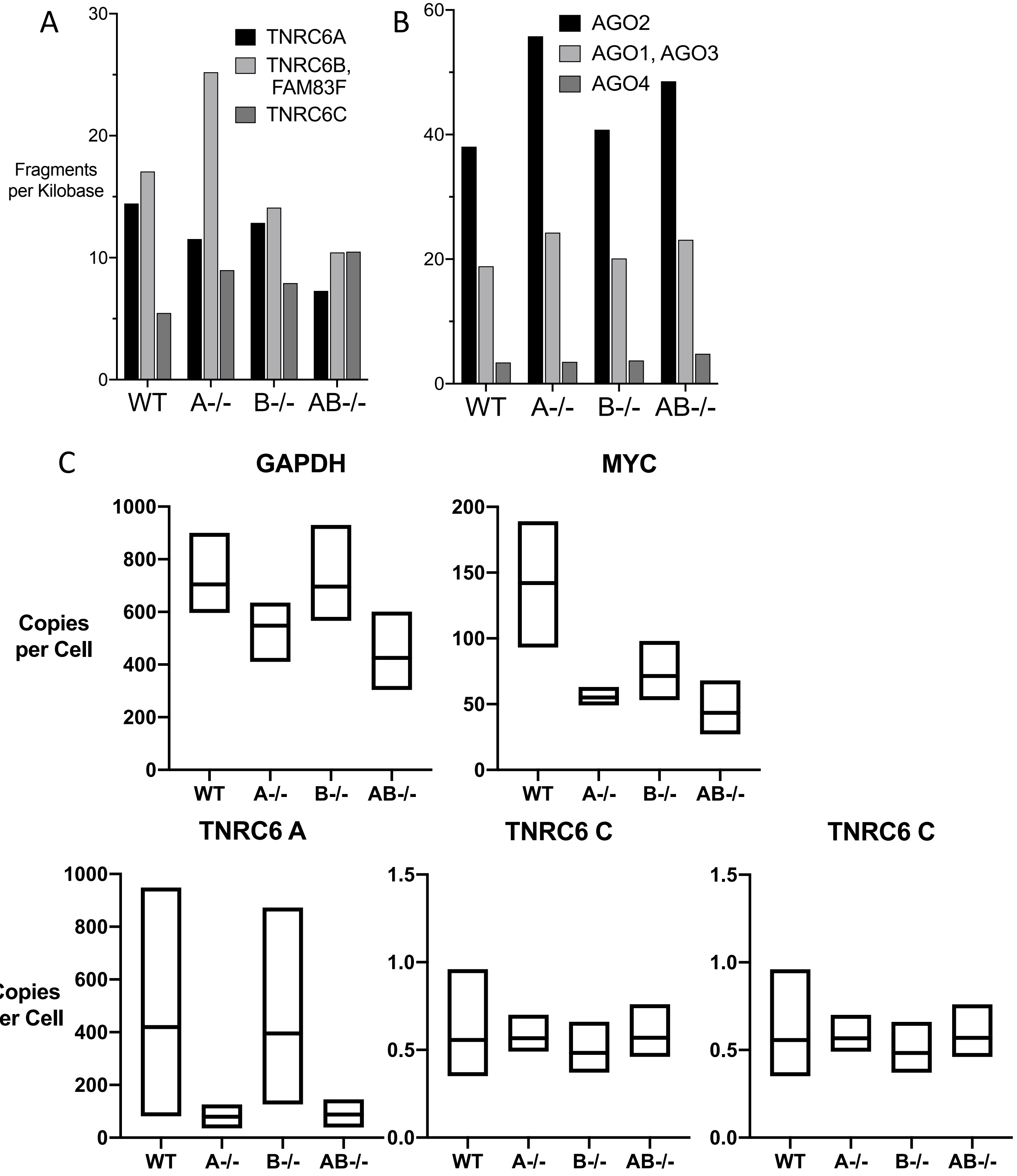


**SUPPLEMENTARY FIGURE 2.** Changes in TNRC6 mRNA and Protein levels after knockout. (A) Bar graph of TNRC6 paralog's fragments per kilobase values obtained from whole cell RNA sequencing of knockout cell lines. TNRC6B and FAM83F are overlapping genes. (B) Bar graph of Ago paralog's fragments per kilobase values obtained from whole cell RNA sequencing of knockout cell lines. AGO1 and AGO3 are overlapping genes. (C) mRNA quantification of the RNA copies per cell of GAPDH, MYC, TNRC6 A, TNRC6 B and TNRC6 C in knockout cell lines.

**SUPPLEMENTARY TABLE 1.** Primers for mRNA quantification and qPCR

| Target | | Primer 5' to 3' |
|---|---|---|
| TNRC6 A fragment for mRNA quantification | FWD1 | ATTTAGGTGACACTATAGAAGACAGGTCCGTTTCCGGTTG |
| | FWD2 | ATTTAGGTGACACTATAGAAGGGGAGACCCTCCAAAGTCTA |
| | REV | ATCATTCCCCTTTCCCTTCTCC |
| TNRC6 B fragment for mRNA quantification | FWD | ATTTAGGTGACACTATAGAA CCTATGAGTATGGCTCCTGTTG |
| | REV | AGGGTTTGTGAGTTCCCTATTT |
| TNRC6 C fragment for mRNA quantification | FWD | ATTTAGGTGACACTATAGAAGAGCAGCAGTCTTCACCCAA |
| | REV | GCAGCGTGGCATTCTGAGAT |
| MYC fragment for mRNA quantification | FWD | ATTTAGGTGACACTATAGAACTTGTACCTGCAGGATCTGA |
| | REV | CTCTTGGCAGCAGGATAGT |
| GAPDH fragment for mRNA quantification | FWD | ATTTAGGTGACACTATAGAACTCTGCTCCTCCTGTTCGAC |
| | REV | TTGATTTTGGAGGGATCTCG |
| TNRC6 A qPCR | FWD | AGCAAGCACAGGTACATCAG |
| | REV | CAGTTGTGGCTGGAGTAGAAG |
| TNRC6 B qPCR | FWD | |
| | REV | |
| TNRC6 C qPCR for knockdown efficiency | FWD | CTGGAGGTCTAAGCATTGGGC |
| | REV | TCAGGGTCATTCTCAGGGTCAA |
| TNRC6 C qPCR for quantification | FWD | |
| | REV | |
| GAPDH qPCR | FWD | GTCAACGGATTTGGTCGTATTG |
| | REV | TGTAGTTGAGGTCAATGAAGGG |
| MYC qPCR | FWD | TCTTCCCCTACCCTCTCAAC |
| | REV | ACCAACCCCAATGTACTTCAG |
| HPRT1 qPCR | FWD | AGTTCTGTGGCCATCTGCTTAGTAG |
| | REV | AAACAACAATCCGCCCAAAGG |

**SUPPLEMENTARY TABLE 2.** siRNAs

| Target | | Primer 5' to 3' |
|---|---|---|
| siGL2 | AS | UCGAAGUAUUCCGCGUACGdTdT |
| | SS | CGUACGCGGAAUACUUCGAdTdT |
| siTNRC6 C Pool | AS1 | AAGUGGACGUUUGUGGUUCdTdT |
| | SS1 | GAACCACAAACGUCCACUU |
| | AS2 | UACUGAUGUCAACCUGGAAGUGUAGAA |
| | SS2 | CUACACUUCCAGGUUGACAUCAGA |
| | AS3 | UGACAUUCAUGUUUGGGUUCAGUCCAG |
| | SS3 | GGACUGAACCCAAACAUGAAUGUCA |

**SUPPLEMENTARY TABLE 3.** PCR primers using in gel electrophoresis and qPCR for alternative splicing analysis

| Application | Gene Name | Target | | Sequence (5'-3') |
|---|---|---|---|---|
| Agarose gel electrophoresis | APIP | E1 | FWD | GCTCGGGAGGGAGACTGTTGTTC |
| | | E3 | REV | CTGAATTCGTTCCTTTTGCACTCC |
| | | 5'UTR | FWD | AAAGCCGTGCGGAGATTGGAGG |
| | | E3/4 | REV | CATGTCTTCAGGCTGAATTCGTTC |
| | EPB41L2 | Ex12 | FWD | AGCATTGCCGTGGTACAAGA |
| | | EX15 | REV | TCGTCTTTCCCAACCTCTGC |
| | FKBP14 | 1 - E2 | FWD | TGGTCAGCCCATTTGGTTTACC |
| | | 2 - E2 | FWD | GCTCATCATTCCTCCTGCTCTGG |
| | | 1 - E4 | REV | TGGAATGATTCATGGGATCTTGGTC |
| | | 2 - E4 | REV | AGAGAGTTTCCAGTCATCATTAAGATCC |
| | KIF21A | E22 | FWD | AGATGCTTTACTAGGCCATGC |
| | | E25 | REV | ATGAGATCTGAAGACAGCGTG |
| | PPIP5K2 | 1 - E25 | FWD | GATAATGATGATGAACCACATACTTC |
| | | 2 - E25 | FWD | AGAGATGAAGTTGATCGAGCTGTG |
| | | 1 - E27 | REV | CTACAGTAGGATTCTGCTTCTGTTCC |
| | | 2 - E27 | REV | CACAAGAGTTCTTGGTGTTCTCAGG |
| | PHLDB1 | 1 - E9 | FWD | AGAAGGAGCAGAAGGCAGTGGATC |
| | | 2 - E9 | FWD | AGAAGCTGGTGGCCTTGGAGACAG |
| | | E10 | REV | TCCAACTGCTGGAACTCCAAGTC |
| | RUBCN | E12 | FWD | AGTTCAGCTCACGTGATTCGGCAC |
| | | E14 | REV | AAGGATTTGCTGCTTGAGGCTGTG |
| | | E11 | FWD | TACCAGGAGGCTGAGCACGGAAGC |
| | | E15 | REV | TCAGCAGACGTGGAGTGCAGGAAG |
| | TBC1D5 | 1 - E3 | FWD | AGTTACTTTTGGTGACGCTGTCC |
| | | 2 - E3 | FWD | ACTAGACATCCTCTGCAGCCAGAAG |
| | | E5 | REV | CTGCTTCTCAGCTGCCCATTAATC |
| | | E5/6 | REV | GAAATAGCTTCCAGCAAATGCTGC |
| QPCR | EPB41L2 | 1 - Ex13 | FWD | GAAGGGGATAATATTTATGTCAGAC |
| | | 1 - Ex14 | REV | TGAGTTCACTAATGCTAGCCTG |
| | | 2 - Ex13 | FWD | AAAATTCCTTGAGAGTAGAAGGG |
| | | 2 - Ex14 | REV | CGGCTCAGGTGTGGATTC |
| | | 3 - Ex13 | FWD | AGAGTAGAAGGGGATAATATTTATGTCAG |
| | | 1 - Ex12/15 | FWD | CAGCTCATTGAAGGAAAGAG |
| | | 1 - Ex15 | REV | ACTCTTTCGTCTTTCCCAACC |
| | | 2 - Ex12/15 | FWD | CAGCTCATTGAAGGAAAGAGT |
| | | 2 - Ex15 | REV | GCTCCACCTTACGAATCTCC |
| | FKBP14 | E3 | FWD | AGCGCATGTCACATCTCTGTCTAG |
| | | E2/4 | FWD | GGAAAAGAAGGAAAAGGTAAAATTCC |
| | | E3 | REV | AGATCTAGACAGAGATGTGACATGC |
| | | E2/4 | REV | CTTTCTGGGGGAATTTTACCTTTTCC |
| | KIF21A | E22-23 | FWD | ACAAGATCTAGATAGCGTACCATTAG |
| | | E24 | REV | TCTGATCCTGGGCTGTTTAAAG |
| | | E22 | FWD | CAGAAATAACCAGTGCTACCCA |
| | | E23-24 | REV | CCTCTACATTTTCTAATGGTACGC |
| | | 1 - E22/24 | FWD | CTAGGCCATGCTTTACAAGAAAATG |
| | | E25 | REV | ATGAGATCTGAAGACAGCGTG |
| | | 2 - E22/24 | FWD | CTAGGCCATGCTTTACAAGAA |
| | | E26 | REV | CTCCTTCGGGCCTTGTTCTT |
| | RUBCN | E13/E14 | FWD | GGCCTCGATGTTCTCAGATGCTGA |
| | | E12/14 | FWD | GAATTTGAAATCCAAGATGCTGACA |
| | | E13/E14 | REV | CCTTCTGATGTCAGCATCTGAGAAC |
| | | E12/14 | REV | CCTTCTGATGTCAGCATCTTGGATT |
